# Nuclear transcriptomes at high resolution using retooled INTACT

**DOI:** 10.1101/180992

**Authors:** Mauricio A. Reynoso, Germain C. Pauluzzi, Kaisa Kajala, Sean Cabanlit, Joel Velasco, Jérémie Bazin, Roger Deal, Neelima R. Sinha, Siobhan M. Brady, Julia Bailey-Serres

**Affiliations:** Center for Plant Cell Biology, Department of Botany and Plant Sciences, University of California, Riverside, CA, 92521; Department of Plant Biology, UC Davis,Davis, CA, 95616; Genome Center, UC Davis, Davis, CA, 95616; IPS2, Institute of Plant Science-Paris Saclay (CNRS-INRA), University of Paris-Saclay, F-911405, Orsay, France; Department of Biology, Emory University, Atlanta, GA, 30322

**Keywords:** Nuclei purification, nuclear transcriptome, rRNA degradation, T-DNA insertion mapping

## Abstract

Isolated nuclei provide access to early steps in gene regulation involving chromatin as well as transcript production and processing. Here we describe transfer of the Isolation of <u>N</u>uclei from <u>TA</u>gged specific <u>C</u>ell <u>T</u>ypes (INTACT) to the monocot rice (*Oryza sativa* L.). The purification of biotinylated nuclei was redesigned by replacing the outer nuclear envelope-targeting domain of the Nuclear Tagging Fusion (NTF) protein with an outer nuclear envelope-anchored domain. This modified NTF was combined with codon optimized *E. coli BirA* in a single T-DNA construct. We also developed inexpensive methods for INTACT, T-DNA insertion mapping and profiling of the complete nuclear transcriptome, including a rRNA degradation procedure that minimizes pre-rRNA transcripts. A high-resolution comparison of nuclear and steady-state poly (A)^+^ transcript populations of seedling root tips confirmed the capture of pre-mRNA and exposed distinctions in diversity and abundance of the nuclear and total transcriptomes. This retooled INTACT can enable high-resolution monitoring of the nuclear transcriptome and chromatin in specific cell-types of rice and other species.

**Summary:** Improved technology and methodology for affinity purification of nuclei and analysis of nuclear transcriptomes, chromatin and other nuclear components.

## Introduction

Since the mid-1980s, gene expression studies in plants that sample steady-state levels of polyadenylated (poly (A)^+^) mRNAs obtained from organs have propelled the understanding of gene regulation and cellular processes. The sampling of poly (A)^+^ mRNAs from laser-captured tissue sections or cells sorted based on expression of a fluorescent marker has further refined knowledge of gene activity, particularly at the cell type-specific level and even more recently at the single cell level (reviewed by (Bailey-Serres, 2013; Efroni et al., 2015). Poly (A)^+^ transcript levels provide a snapshot of the output of transcription and mRNA stability in the whole cell. RNA-sequencing (RNAseq) on cell type-specific transcriptomes additionally yields information about the sites where transcription initiates and polyadenylation occurs, and provides insights into the alternative splicing and retention of introns. However, steady-state transcript abundance may not accurately predict nascent transcription or the transcripts undergoing translation, particularly as plants modulate these processes under environmental stress (Park et al., 2012). Early analyses of plant transcript dynamics that used hybridization kinetics gleaned that nuclear and polyribosomal (polysomal) mRNA populations are highly distinct in tobacco (*Nicotiana tobaccum*) leaves (Goldberg et al., 1978). Others have compared nuclear poly (A)^+^ (Zhang et al., 2008) and total nuclear RNA (Deal and Henikoff, 2010) to total poly (A)^+^ mRNA of *Arabidopsis thaliana* using microarrays. More recently, two studies have used nuclear RNAs to evaluate their structure and RNA-protein interactions (Gosai et al., 2015; Foley et al., 2017). To date, there has been no comparison of nuclear RNA with total cellular poly (A)^+^ RNA using high-throughput RNA-seq for any plant. Technologies that facilitate these comparisons in model and crop plants could aid the study of regulatory processes that are obscured by exclusive sampling the total poly (A)^+^ mRNA. Genome-scale sampling of nuclear RNA can be accomplished by isolation of nuclei using differential centrifugation, <u>F</u>luorescence <u>A</u>ctivated <u>N</u>uclear <u>S</u>orting (FANS) (Zhang et al., 2008) or Isolation of <u>N</u>uclei <u>TA</u>gged in specific <u>C</u>ell <u>T</u>ypes (INTACT) (Deal and Henikoff, 2010; Deal and Henikoff, 2011; Bailey-Serres, 2013). The latter two techniques were established for *A. thaliana* and take advantage of transgenes that encode green fluorescent protein (GFP) fused to the core nucleosome protein histone 2A or a biotinylated chimeric fusion protein with affinity to an outer nuclear membrane (ONM) protein, respectively. FANS requires preparation of nuclei from fresh tissue and the use of a fluorescent activated cell sorter. INTACT, on the other hand, is an inexpensive affinity purification method that is routinely performed on frozen tissues, enabling studies of rapid changes in gene regulation. Because INTACT entails the use of a transgene that can be driven by spatially-or temporally-regulated promoters, it enables sampling of nuclei of subpopulations of cell types or tissues within intact organs (Deal and Henikoff, 2010).

INTACT provides access to nuclei, including DNA, histones, other nuclear proteins, pre-ribosomal RNAs, pre-mRNA and other RNAs within the nucleoplasm. There are two required components for this technology. The first is a synthetic Nuclear Tagging Fusion (NTF) protein comprised of an ONM-targeting domain (Trp–Pro-Pro [WPP]) from *RAN GTPASE-ACTIVATING PROTEIN 1* (*RanGAP1*) (Rose and Meier, 2001), fused to GFP and a biotin ligase target peptide (BLRP). The second is an *E. coli* Biotin Ligase (*BirA*) that is expressed in the same tissue (Deal and Henikoff, 2010); (Deal and Henikoff, 2011). The WPP domain interacts with a coil-coiled domain of a WPP-interacting protein (WIP). WIP itself is anchored via a C-terminal transmembrane domain or KASH (Klarsicht / ANC–1 / Syne-1 Homology) to the ONM and interacts with other proteins of the inner nuclear membrane (reviewed by Zhou et al., 2015; Rose and Meier, 2001; Xu et al., 2007). The interaction between the coil-coiled domain of WIP and the WPP domain places the NTF on the exterior of the nucleus. The biotinylation of the NTF enables the use of streptavidin-coated magnetic beads to selectively purify tagged nuclei from crude extracts of plant tissues. As WIP is directly anchored to the ONM and WPP is not, we considered that replacing the WPP domain with a KASH domain would provide more stable anchoring of the NTF to nuclei.

We recently translated INTACT to tomato (*Solanum lycopersicum*) (Ron et al., 2014). Here, we retooled both the NTF and *BirA* for efficient INTACT in monocots, specifically rice. We also refined inexpensive methods for preparation of RNA-seq libraries from the complete nuclear transcriptome by including subtraction of pre-rRNA. To demonstrate one use of this technology, we used an INTACT construct driven by the near-constitutive CaMV 35S promoter (*p35S*) to isolate nuclei and performed a comparative RNA-seq analysis on nuclear and total poly (A)^+^ mRNA from root tips. The results demonstrate notable distinctions between the gene transcripts represented and their abundance in the nuclear and poly (A)^+^ transcriptomes. We discuss applications of INTACT to the study of nuclear processes of gene regulation, from chromatin to mRNA export, and at the organ to cell-specific level.

## RESULTS AND DISCUSSION

### Generation of transgenic rice lines for purification of nuclei using INTACT

The retooling of INTACT for rice included modifications to improve biotinylation and to establish a stable interaction with the ONM. First, due to the greater GC content of the third codon position of genes in rice as compared to *A. thaliana* (Tatarinova et al., 2013), we synthesized a rice codon usage-optimized version of *E. coli BirA*. This was included in a binary T-DNA vector along with the Nuclear Tagging Fusion (NTF) construct. In *A. thaliana* and *S. lycopersicum,* to establish a tagged nucleus, the NTF used was the WPP of RanGAP1 (Deal and Henikoff, 2010; Ron et al., 2014). The rice *ACTIN2* (*ACT2*; *LOC_Os10g36650*) promoter was selected to drive expression of *BirA* as its transcript is detected across cell types based on available expression data (Jain et al., 2007; Jiao et al., 2009). Because WPP association with the ONM is developmentally regulated (Xu et al., 2007; Zhao et al., 2008), we tested whether the C-terminal region of OsWIP (c-OsWIP, including a coil-coiled and a C-terminal transmembrane domain) (reviewed by (Zhou et al., 2015)) could be used for INTACT. We designed three NTF constructs using rice WPP and c-OsWIPs domains for evaluation. NTF1 combined the OsWPP domain of the *RanGAP1* ortholog (*LOC_Os05g46560.1*; (Rose and Meier, 2001)) with GFP and a C-terminal BLRP domain (Fig. 1A). For NTF2 and NTF3, we replaced the WPP domain with the c-OsWIP from two rice orthologs of *Arabidopsis* WIP1: *LOC_Os09g30350.1* (Os*WIP2*) and *LOC_Os08g38890.1* (*OsWIP3*) (Xu et al., 2007). To position this domain in the ONM so that the GFP and BLRP of the NTF would be on the cytoplasmic face of the nucleus, we placed the BLRP at the N-terminus, fused to GFP and then followed by the c-OsWIP (Fig. 1A).

**Figure 1.**
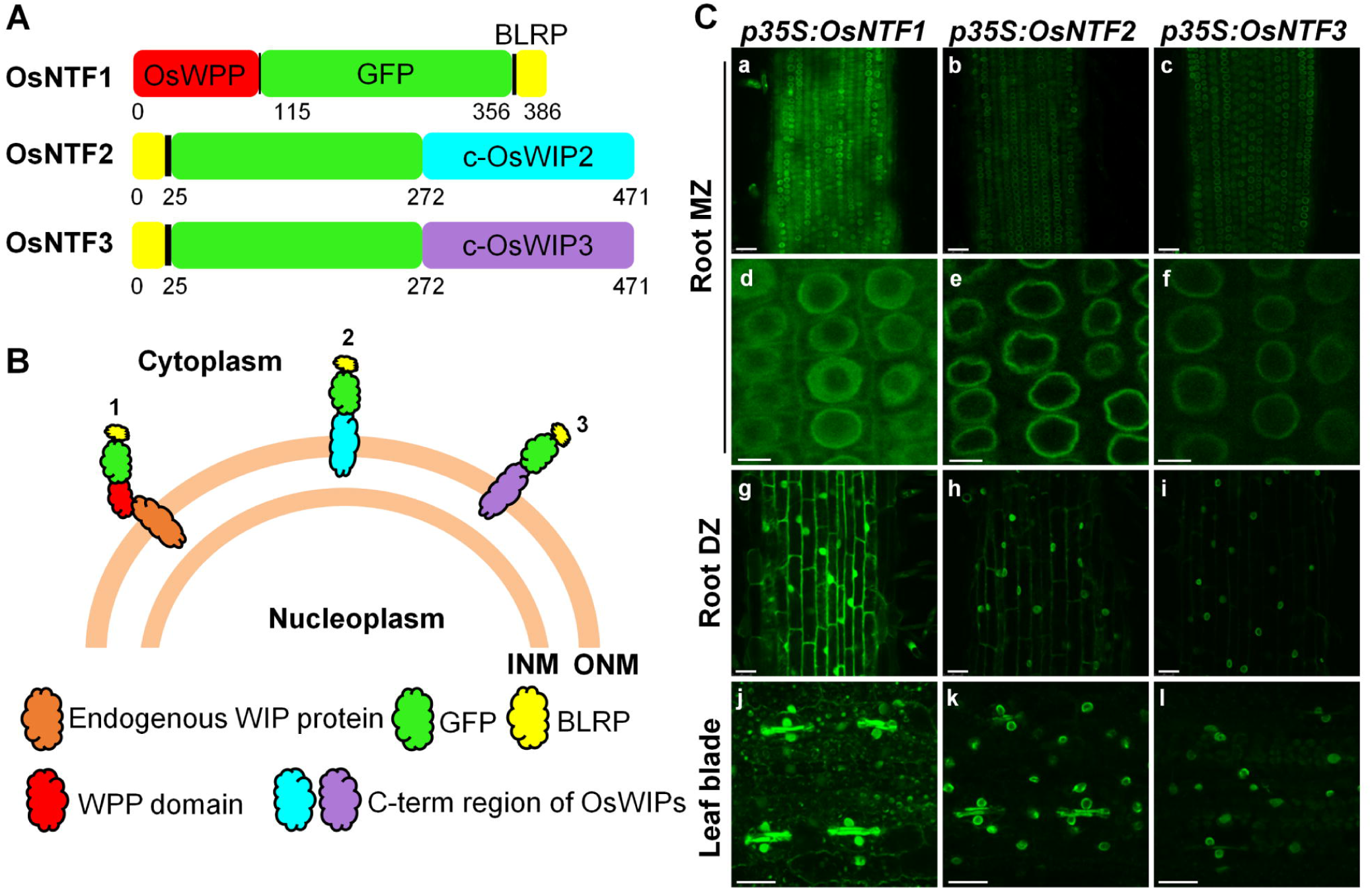
Three Nuclear Tagging Fusion (NTF) proteins and their subcellular localization. **A,** Three synthetic NTF proteins were tested. Each was composed of three functional regions: a Biotin Ligase Receptor Peptide for biotinylation (BLRP), a Green Fluorescent Protein (GFP) for visualization, and a WPP domain or a C-terminal region of WIP proteins for targeting to or anchoring on cytoplasmic side of the nuclear envelope, respectively. The nuclear targeting domains are *OsWPP* (1), c-OsWIP2 (2) and c-OsWIP3 (3). **B,** Diagrammatic representation of predicted NTF location in the nuclear-envelope-cytoplasm interface. *OsNTF*1 is associated with the nuclear envelope via interaction between its WPP domain and endogenous WIP proteins. The C-terminal region of WIP of*OsNTF2* and *OsNTF3* may be sufficient for integration into the outer nuclear envelope. **C,** Representative confocal images of GFP fluorescence in organs from 7-day-old plants grown in a growth chamber expressing a *p35S:0sNTFs.* a to f: root meristematic zone (Root MZ); g to i: root differentiated zone (Root DZ) 1 cm from the root tip, epidermal cells; j to 1: blade region between the midvein and outer margin of a newly expanding leaf blade, guard cells and epidermal cells are visible. Leaves are maximum intensity Z-projections of a confocal Z series. INM: Inner Nuclear Membrane, ONM: Outer Nuclear Membrane. Scale bars are: a-c, g-l: 25 μ M, d-f: 5μ M.

Independent transgenic rice lines were produced that express each NTF version under the control of the *p35S* (*p35S:OsNTF*s). There was no detectable phenotypic consequence of expression of any of these constructs on development or fertility under standard culture or greenhouse conditions. The sites of T-DNA insertion in 12 independent transgenics were determined using a high-throughput method that captures T-DNA border regions and flanking DNA by hybridization to biotinylated probes (Supplemental Table S1; Supplemental Results). Sites of insertion were validated by PCR. The *p35S:Os*NTF*1*, *p35S:OsNTF2* and *p35S:OsNTF3* lines used for subsequent analyses had moderate level of GFP fluorescence and a single insertion in chromosome 6, 1 and 1, respectively.

### Detection of nuclear envelope-associated and anchored biotinylated-GFP

Confocal microscopy was used to monitor the cellular distribution of the three *Os*NTF proteins in cells of roots and leaves of 7–day-old seedlings. *Os*NTF1 is predicted to associate with the ONM via protein-protein interaction with a WIP (Fig. 1B). Most of the GFP signal from *Os*NTF1 was localized at the periphery of nuclei in roots (meristematic and differentiated region) and expanding juvenile leaf blades (Fig. 1C; Fig. S1). The GFP signal was also abundant in the cytoplasm of the root meristematic zone as well as the differentiated zone (epidermal cells), where it was displaced by the central vacuole. *Os*NTF2 and *Os*NTF3 were designed to be anchored to the ONM via the C-terminal transmembrane domain of WIP (Fig 1B). The sharper halo of GFP signal around the nuclei in *p35S:OsNTF2 -* and *p35S:OsNTF3*-expressing plants indicates a more integral association of WIP than WPP with the nuclear membrane (Fig 1C). *p35S:OsNTF2* and *p35S:OsNTF*3 plants had less GFP signal in the cytoplasm than *p35S:OsNTF1* plants, even though the highly-expressed *p35S* was used for all constructs. Lower cytoplasmic accumulation of NTF2 and NTF3 is quite evident in the guard cells of the leaf blade. The high cytoplasmic GFP signal in multiple *p35S:OsNTF1* lines (Fig. S1), suggest that the amount of protein produced exceeded available WIP or other nuclear anchoring proteins. Alternatively, *Os*NTF2 or *Os*NTF3 protein that does not integrate into the ONM may be more readily degraded than *Os*NTF1.

To confirm that *Os*NTFs were biotinylated *in planta*, root tissue was solubilized in an SDS-containing buffer and proteins were fractionated by SDS-polyacrylamide gel electrophoresis (SDS-PAGE), blotted to a membrane and probed with streptavidin-conjugated to horseradish peroxidase, which binds with high affinity to biotin and enables enzymatic detection, respectively. Biotinylated proteins with the expected electrophoretic mobility were detected in *p35S:OsNTF1* -, *2*-and *3*-expressing plants (Fig. S2). These results demonstrate that a chimeric GFP can associate with or be anchored to the ONM in rice, supporting the use of these *Os*NTF-expressing lines for affinity purification of nuclei.

### *Os*NTFs with a WPP or KASH domain enable rapid purification of nuclei

Nuclei from frozen tissue of 7–d-old seedlings expressing *Os*NTFs were purified by INTACT following the procedure described for *A. thaliana* (Deal and Henikoff, 2010), with minor modifications (Fig. 2A; See Methods). After affinity capture by binding to an excess of streptavidin-coated magnetic beads and extensive washing, the nuclei were stained with 4’, 6–diamidino–2-phenylindole (DAPI) to detect nucleic acids by fluorescent microscopy (Fig. 2B). Nuclei were purified with all three *Os*NTF versions in significant excess over nuclei captured by the same procedure from roots and shoots of non-transgenic seedlings (Fig. 2C and D; Fig. S3A). To further evaluate INTACT with these constructs, we examined the presence of chromatin by monitoring histone H3 abundance. Histone H3 levels were similar for all four genotypes in the input (Fig. 2E, F), whereas H3 levels captured by INTACT were higher from *p35S:OsNTF2* and *p35S:OsNTF3* tissue in accordance with greater number of nuclei purified (Fig. 2C and 2D). The same samples in panels C to F, indicate that histone H3 levels were efficiently purified from *Os*NTF2 and *Os*NTF3 seedlings. Thus, the INTACT constructs with a KASH domain display a more defined nuclear association, less cytoplasmic signal, and were suitable for capture of nuclei and associated chromatin from seedling tissues. There was some variation in nuclear yield between NTF constructs, insertion events, organ sampled and experiments. For example, nuclear yields were higher in roots than shoots for OsNTF3 but not OsNTF2 (Fig. 2C and 2D) and high OsNTF1 accumulation as in the line presented in Fig S1, provided similar levels of purified nuclei as *Os*NTF2 and *Os*NTF3 lines (Fig. S3A). We conclude that all three NTF constructs prepared for and expressed in rice worked for the intended purpose. A *p35S:OsNTF2* line that consistently gave good yields of nuclei from seedling and field grown tissue was used for further evaluation of INTACT purified protein and RNA.

**Figure 2.**
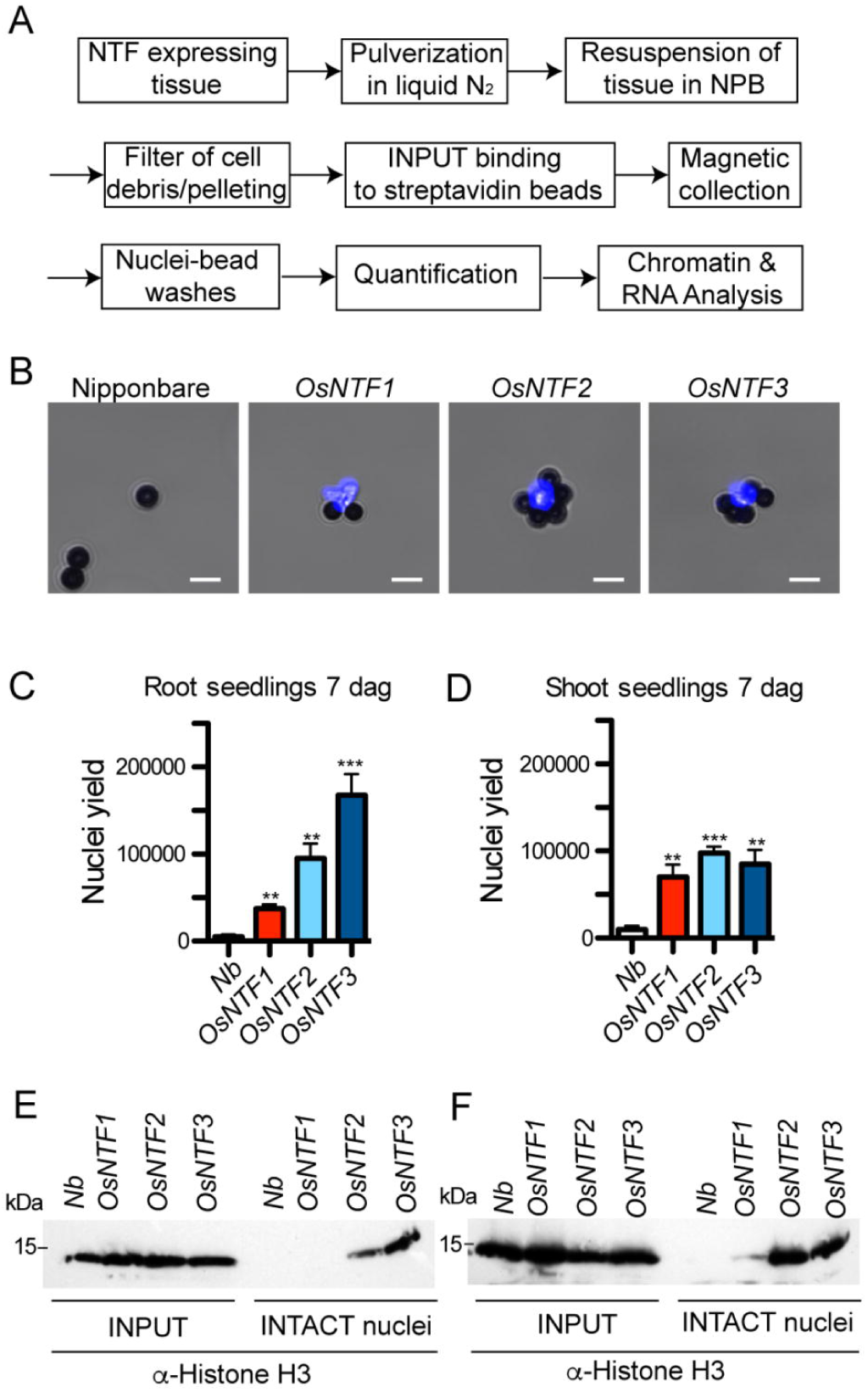
Nuclei isolation using INTACT. **A,** Scheme of INTACT purification. **B,** Capture of biotinylated nuclei from roots of transgenic lines carrying *p35S:OsNTFI*, 2 or 3 *andpACT2-BirA* transgenes by use of streptavidin-coated magnetic beads. Nuclei were stained with DAPT to visualize DNA. Viewed with mixed white light and DAPI-channel fluorescence illumination. Nuclei (bright blue) surrounded by 2.8 flm spherical beads (dark grey) are visible. Scale bar is 5 μm. **C** and **D,** Nuclei were purified from roots and shoots of 7-d-old seedlings expressing the different OsNTF versions, stained with propidium iodide for visualization and counted by use of an hemocytometer. Asterisks indicate significant differences in an unpaired two-tailed Student’s t-test (** p<0.01 and *** p<0.001) in a comparison to nuclei purified with non-transgenic Nipponbare (Nb). **E** and **F,** Detection of Histone 3 (H3) protein in nuclei before (crude nuclei) or after purification by INTACT (INTACT nuclei). For each sample, an equal amount of starting material (0.5 g) was used to obtain nuclei and a proportional amount of each yield was solubilized in an SDS-containing buffer and fractionated by electrophoresis in an 12% (w/v) polyacrylamide-SDS gel, transferred to a membrane and processed with commercial monoclonal antibody against Histone 3 (H3) (α-Histone H3). Molecular weight marker shown on the left. The expected size of Histone H3 is 15.27 kDa.

To compare and contrast the nuclei obtained by INTACT or conventional centrifugation methods we monitored abundant proteins. First, histone H3 protein levels of INTACT and sedimented nuclei, of shoot tissue of 5–week-old plants grown under field conditions were compared. Total cellular and nuclear proteins were fractionated by SDS-PAGE and silver stained (Fig. 3A). As expected, a greater number of protein bands were detected in the total protein extract than in isolated nuclei. Both nuclear preparations were enriched in low-molecular mass proteins of 9.3 to 19.7 kDa apparent mass, characteristic of the abundant core histones. INTACT nuclei showed an enrichment of histone H3 as compared to the conventionally isolated nuclei (Fig. 3B). The abundance of ribosomal subunit protein S6 (RPS6) was highest in the crude extract and less abundant in INTACT nuclei (Fig. 3C). We anticipated that RPS6 would also be detected in the nuclear fraction due to ribosome biogenesis in the nucleolus, but the level was far less than in the total extract indicating that nuclear pre-ribosomes are highly outnumbered by cytoplasmic ribosomes. The higher level of RPS6 relative to histone H3 in the conventionally purified nuclei could reflect either chromatin loss, cytoplasmic contamination, or better maintenance of the nuclear membrane-rough endoplasmic reticulum connection. Altogether, these results confirm the effective purification of chromatin by INTACT with *Os*NTF2 in rice.

**Figure 3.**
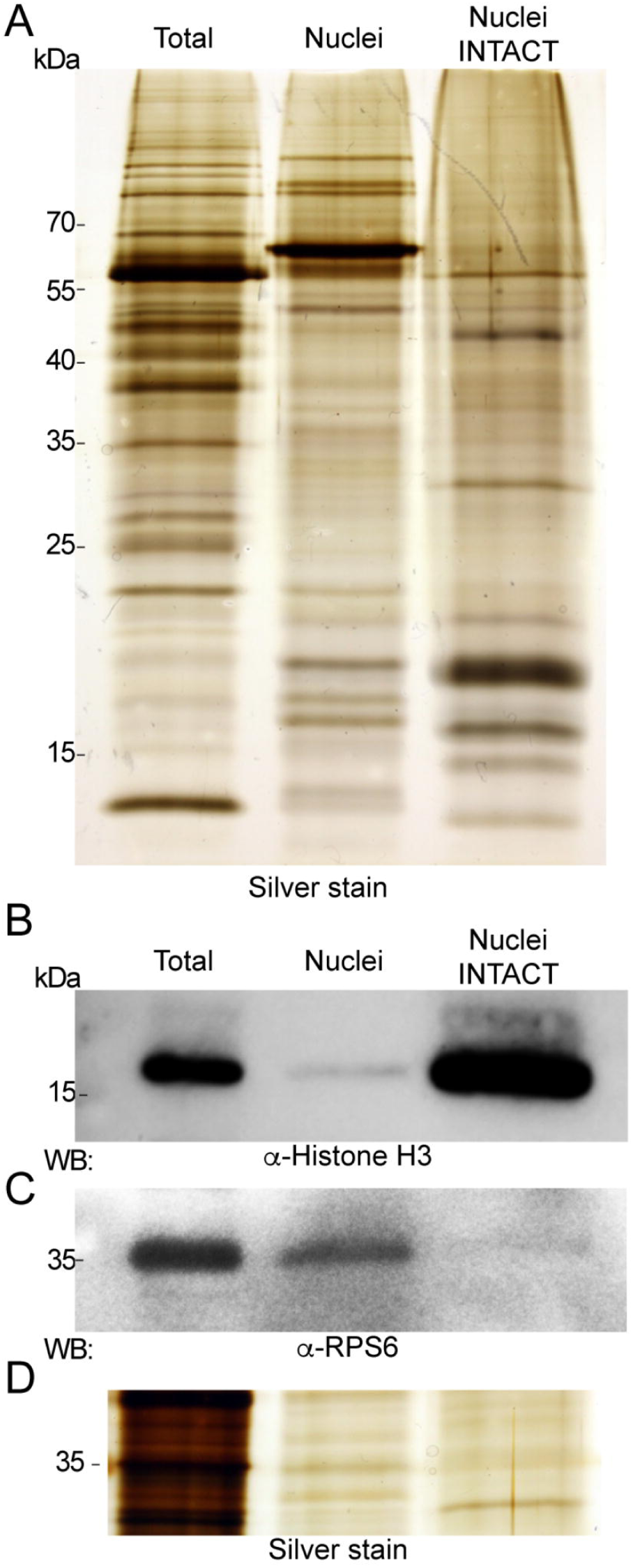
Evaluation of protein composition from INTACT-captured nuclei. **A,** Proteins isolated from 5-wk-old field grown plant (Total), nuclei prepared by conventional sedimentation methods (Nuclei) and using INTACT (Nuclei INTACT) were separated by 12% (w/v) SDS-PAGE and visualized by silver staining. **B,** Total and nuclear proteins were analyzed by western blot using α-Histone H3 and **C,** ribosomal protein α-RPS6 antibodies or silver stained (lower panel). Molecular weight markers are shown on the left. **D,** Protein loading is the same for panels B, C and D, based on protein quantification (see methods). Protein loading in panel A was adjusted to optimize the visual comparison between samples.

### Preparation of nuclear RNA libraries with pre / rRNA subtraction

Next we explored the use of INTACT to evaluate the complete nuclear transcriptome by RNA-seq using root tips of 7-d-old *p35S:NTF2* seedlings. To selectively remove the abundant pre-and processed rRNAs produced by the 45S rDNA and 5S rRNA clusters (Fig. 4A), we developed method to remove rRNA that did not rely on a commercial kit. Nuclear RNA was hybridized to DNA oligos complementary to 25S, 18S, 5.8S and 5S rRNAs and the DNA: RNA hybrids were selectively degraded at high temperature with a thermostable RNase H. RNA-seq run confirmed this rRNA degradation method reduced reads corresponding to mature rRNAs but not the transcribed spacer sequences of preRNA (Fig. 4Ai). As DNA oligos corresponding to the pre-rRNA external transcribed spacers (5’ ETS and 3’ ETS) and internal transcribed spacers (ITS1 and ITS2) were not included in the initial rRNA degradation assay, the presence of the 5’ETS, ITS1, ITS2 and 3’ETS regions confirmed that nuclear RNA had been isolated. We subsequently adjusted our method to include DNA oligos to subtract mature rRNA as well as pre-rRNA ETS, ITS1 and ITS2 regions (pre-rRNA degraded). This reduced the unprocessed pre-rRNA reads to below the level of rRNA reads in oligo–dT-selected RNA (Fig. 4 Aii-iii). As a control we show that relative coverage over *MALATE DEHYDROGENASE* gene is unaffected (Fig. 4B)

**Figure 4.**
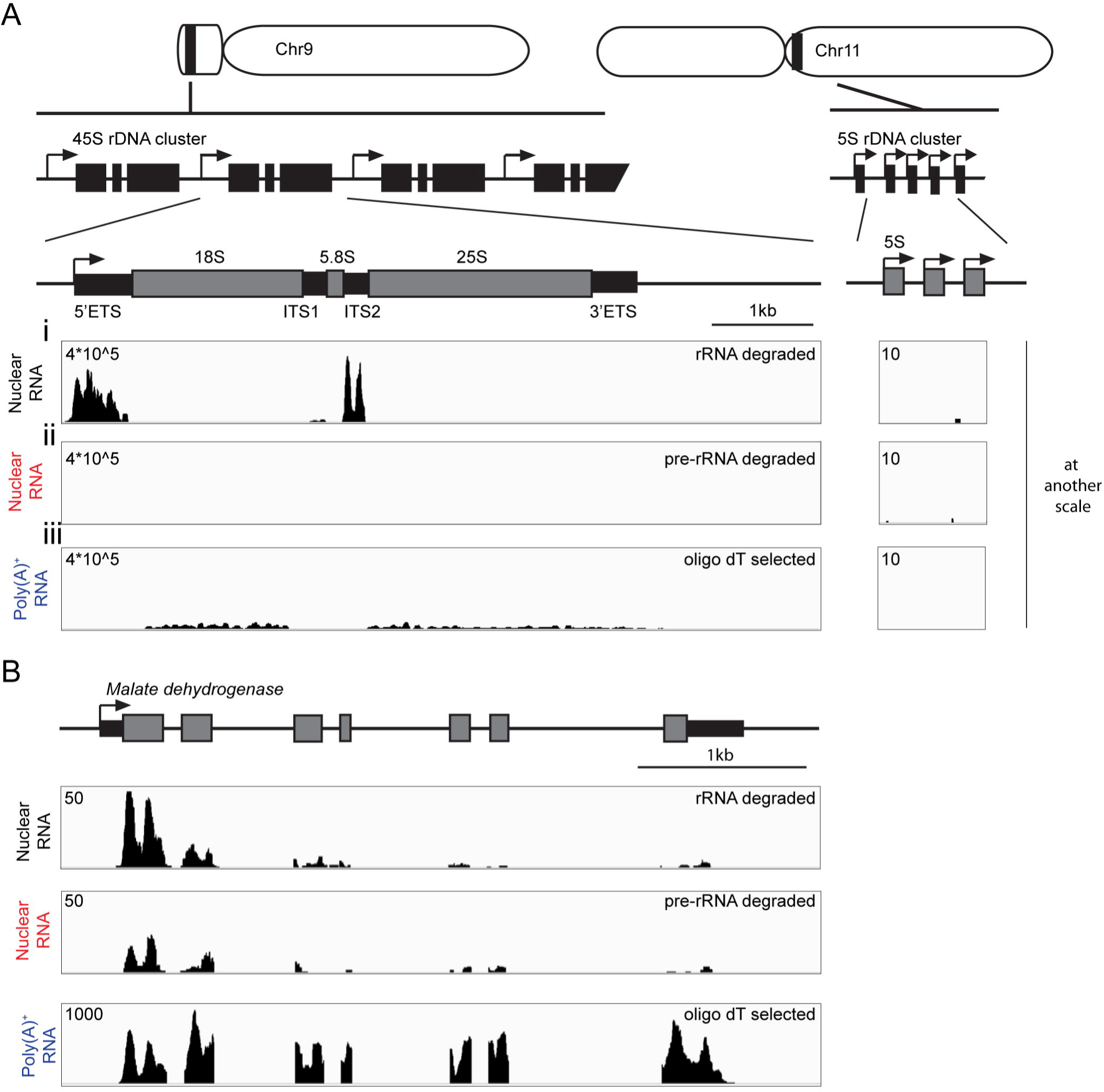
Nuclear transcriptome libraries generated including an rRNA degradation step. **A,** Location, organization of ribosomal DNA (rDNA) and coverage of nuclear transcriptome. Rice 45S rDNA cluster is located in the short arm ofchromosome (Chr) 9 and consists oftandem repeats of 5’ external transcribed spacer (ETS), 18S rRNA, internal transcribed spacer 1 (ITS 1), 5.8S rRNA, ITS2, 25S rRNA and 3’ ETS. 5S rDNA cluster is located in Chr 11 composed of repeats of 5S rRNA. Panels i and ii show read coverage on a 45S and 5S region for nuclear RNA treated with mature rRNA complementary probes **(i)** or full pre-rRNA probes **(ii)**. Panel iii shows the coverage obtained from poly(A)+ selected libraries. **B,** Comparison of read coverage in the nuclear and poly(A)+ RNA transcriptomes for the gene *MALATE DEHYDROGENASE (LOC_OsOIg46070).*

A systematic comparison was made of nuclear RNA and poly(A)^+^ RNA from root tips of seedlings. Nuclei were isolated using *p35S:NTF2* from four biological replicate samples of 200 root tips and used to generate nuclear RNAseq libraries by rRNA subtraction and random–primer-enabled cDNA synthesis (Townsley et al., 2014). Total RNAseq libraries were constructed with oligo dT-selected RNA. After recognizing that the sequences included some reads in very high abundance that were of organellar origin or the ETS, ITS1 and ITS2 regions, in cases where oligos to these regions were not used, we established a pipeline for analysis that commenced with a filtration step. Sequence reads that mapped to mitochondria and chloroplast genomes were removed and the remaining reads were mapped to exons with ≤ 2 mismatches to the Nipponbare genome (IRGSP-1.0.30). This yielded an average coverage of 26.7 (total [poly(A)^+^] RNA) and 1.52 million (nuclear RNA) (Supplemental Table S2a). Despite the difference in read number, subsampling of the libraries confirmed that each library had reached a saturation of transcripts identified as present in each population (Fig S4). Based on reads that mapped to exons, both the poly(A)^+^ RNA libraries (r^2^ ≥ 0.978, Pearson correlation) and nuclear RNA (r^2^ ≥ 0.965) yielded highly consistent results, although there was somewhat greater variation in the nuclear RNA libraries (Supplemental Table S2a).

### The root tip nuclear and poly(A)^+^ RNA transcriptomes differ in complexity and transcript enrichment

To compare the nuclear and poly(A)^+^ transcriptomes we determined the number of genes detected (complexity) and evaluated differences in transcript abundance using averaged data from the four biological replicate samples for each RNA population. First, we evaluated the portion of the transcript mapping to introns, coding regions (CDS) and untranslated regions (UTRs) (Fig. S5). The proportion of intronic to exonic reads (CDS, 5’UTR, 3’UTR) was 2-fold higher in the nuclear RNA in all of the biological replicates. To estimate complexity (*i.e*., the total number of gene transcripts detected), we applied a threshold of > 5 reads per kilobase per million reads (rpkM) on exonic regions (Supplemental Table S2b). In the nuclear RNA ∽18,100 and in the poly(A)^+^ RNA 16,500 protein-coding transcripts were detected, respectively, with 14,264 shared by both populations (Fig. 5A). Gene Ontology (GO) enrichment analysis of the RNAs detected only above the 5 rpkM threshold in the nuclear RNA included meiotic division (BP:adj. P value 0.00448), ribonucleoside binding (MF:0.0039) and peptide receptor activity (MF:0.0043), whereas those detected only in poly(A)^+^ RNA above the 5 rpkM threshold identified phospholipid biosynthetic process (BP: 0.00606) among others (Supplemental Table S2c). This indicates that these mRNAs were highly enriched in the nucleus as compared to the poly(A)^+^ RNA pool, demonstrating that nuclear RNA and poly(A)^+^ measures of transcript abundance yield distinct results. The difference between these read-outs was also evident from a weak positive correlation in abundance of RNAs in the two populations (r 0.298, Pearson correlation) (Fig. 5A). We determined that this was due to both distinctions in transcriptome complexity and transcript abundance. Of the 14,264 transcripts above the 5 rpkM threshold in both populations, 6,234 were significantly enriched in one or the other population (nuclear, 3,152; poly(A)^+^, 3,082; |log_2_ fold difference| > 1, FDR ≤ 0.01) (Supplemental Table S2d). Higher nuclear abundance was found for genes associated with transferase activity (MF:2.87E-5) and cell wall organization or biogenesis (BP:0.0005), whereas higher poly(A)^+^ enrichment was noted for genes associated with translation (BP:1.25E-67) and intracellular transport (BP:1.18E-17) (Supplemental Table S2e).

**Figure 5.**
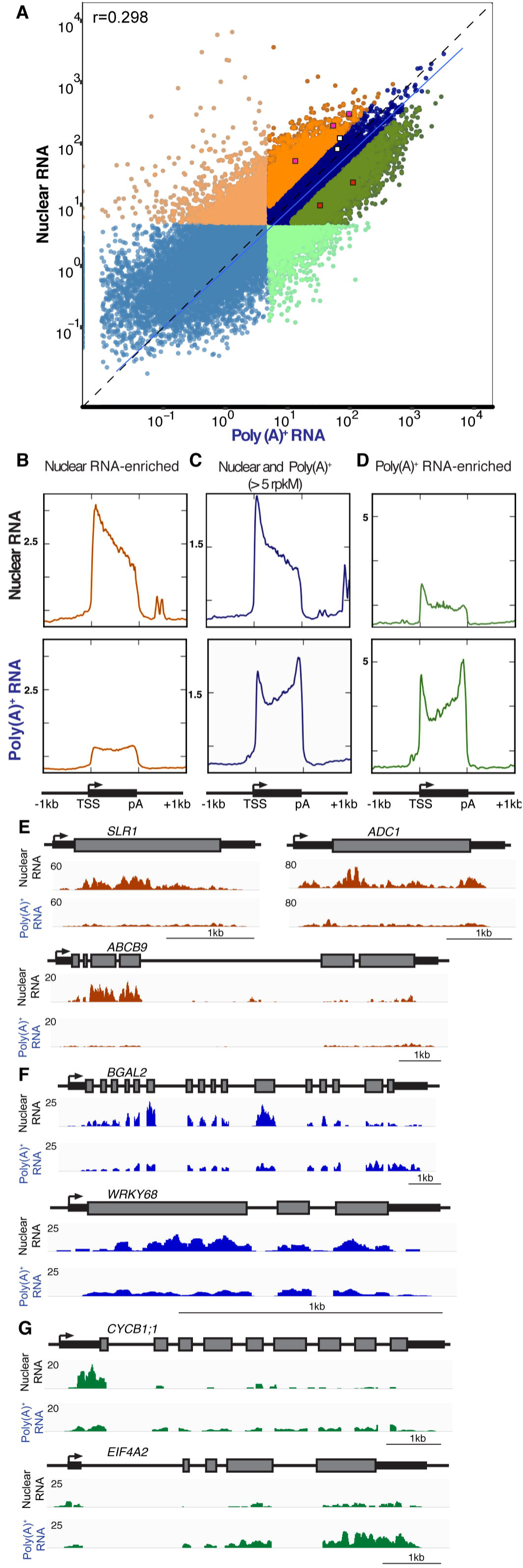
Comparison of the nuclear and total poly(A)+ transcriptome of root tips. Nuclear RNA was extracted from INTACTpurified root tip nuclei and total poly(A)+ RNA was captured using a biotinylated oligo dT. **A,** Scatter plot representing the mean rpkM values on rice annotated genes from four biological replicates for nuclear and poly(A)+ RNA. Light orange and light green dots indicate 3,831 and 2,215 transcripts with >5 rpkM only in nuclear or poly(A)+ RNA, respectively. More intense colored dots indicate transcripts with >5 rpkM in both RNA populations. Dark orange dots indicate 3,152 transcripts differentially enriched in nuclear RNA whereas 3,082 dark green dots represent transcripts enriched in poly(A)+ |log2 FC>I; FDR<0.01|- A linear model is indicated with a solid blue line presenting a 0.298 Pearson correlation value and y=x is indicated by a dashed line. Purple, red and white squares are genes evaluated in panels **E-G. B, C and D,** Nuclear and poly(A)+ RNA coverage evaluation across transcription units. The x-axis represents a transcription unit indicating the transcription start site (TSS), the cleavage and polyadenylation site (pA) and 1 kb region upstream or downstream. The y-axis indicates the relative rt **rt rt** transcript coverage, using different scales for each panel. **B,** nuclear RNA-enriched transcripts from panel A (dark orange dots). **C,** 14,262 transcripts detected in both nuclear and populations at > 5 rpkM. **D**, poly(A)+-enriched transcripts (dark green dots). **E, F and G,** Examples of coverage over individual transcripts. **E**, Nuclear enriched transcripts showing the presence of 5’ bias in nuclear coverage include a *DELLA PROTEIN SLRl (SLRl, LOC_Os03g49990), ARGININE DECARBOXYLASE (ADCl, LOC*_*Os06g04070), ABC TRANSPORTER B FAMILY MEMBER* 9 *(ABCB9, LOC*_*Os02g09720).* These transcripts are indicated in Panel A as purple squares. **F**, Examples of transcripts detected in both nuclear and poly(A)+ include: BETA-GALACTOSIDASE *2 (BGAL2, LOC_Os06g37560)* and *TRANSCRIPTION FACTOR WRKY68 (WRKY68, LOC*_Os*04g51*5*60).* These transcripts are indicated in Panel A as white squares. **G**, Examples ofpoly(A)+ enriched transcripts showing a high 5’ bias in nuclear coverage include a *CYCLINBl-l (CYCRl;l, LOC_OsOlg59l20)* and *EUKARYOTIC INITIATION FACTOR 4A-2 (EIF4A2, LOC_Os02g05330).* These transcripts are indicated in Panel A as red squares

Technical and biological factors could contribute to the lack of congruence between the nuclear and poly(A)^+^ RNA transcriptomes measured by our approach. First, the NTF construct is designed to capture nuclei from cells expressing the 35S promoter, whereas poly(A)^+^ purification accesses all cells following disruption with a chaotropic agent. Therefore, some poly(A)^+^-enriched mRNAs could be those from cells that do not express the *p35S*. Although not exhaustively evaluated, the NTF GFP signal was detectable across cell types in the rice root (Fig. 1C). Second, the greater complexity of the nuclear transcriptome may reflect the absence of the requirement for polyadenylation, thereby including protein-coding RNAs at various points in transcription, processing, nuclear export, and splicing. Indeed, as mentioned, the nuclear RNA samples had greater proportions of introns and 5’UTRs (flanking regions) (Fig. S5).

To further explore if the nuclear RNA pool was enriched for nascent transcripts we determined the average distribution of RNA-seq reads across all individual transcripts detected in nuclear and poly(A)^+^ RNA with >5 rpkM, from the annotated start site (TSS) to the cleavage / poly(A) addition site. This illuminated a bias in RNA-seq reads near the 5’ end of nuclear transcripts in contrast to the more equally distributed reads across the entire transcription unit in poly(A)^+^ RNA (Fig. 5C). The read coverage averaged for all poly(A)^+^ RNAs uncovered a modest 5’-end bias and a more pronounced 3’-end bias. Genome browser views illustrate the average transcript read distribution for *BETA-GALACTOSIDASE 2* (*BGAL2*) and *TRANSCRIPTION FACTOR WRKY68* (*WRKY68*) (Fig. 5A, F). Although these mRNAs have introns, few nuclear RNA reads map to intervening regions, suggesting that splicing occurs with fidelity.

To gain an understanding of the biological basis for nuclear-enriched transcripts, read coverage across the transcription unit was averaged for the RNAs that were significantly enriched in the nuclear or poly(A)^+^ populations. Intriguingly, the nuclear 5’-end bias was less prominent in transcripts with the highest nuclear abundance (Fig. 5B). By contrast, for these RNAs the coverage for the polyadenylated transcript pool was more evenly distributed along the transcription unit. Three nucleus-enriched transcripts illustrate this read mapping pattern in genome browser views: DELLA protein *SLENDER LEAF RICE 1* (*SLR1*), *ARGININE DECARBOXYLASE* (*ADC1*), and *ABC TRANSPORTER B FAMILY MEMBER 9* (*ABCB9*) (Fig. 5A, E). This indicates that export of some nuclear transcripts may be rate limited after completion of splicing and processing. Contrastingly, transcripts with significantly enriched poly(A)^+^ abundance had a 5’-end bias in the nuclear but not poly(A)^+^ read coverage, similar to the average transcript read coverage (Fig. 5D). These transcripts are present at higher levels in the poly(A)^+^ mRNA transcriptome. Genome browser views of *CYCLIN B1-1* (*CYCB1; 1*) and *EUKARYOTIC INITIATION FACTOR 4A-2* (*EIF4A-2*) illustrate a pattern of nuclear depletion relative to poly(A)^+^ transcript accumulation (Fig. 5A, G).

The distinct read coverage patterns between transcripts in the INTACT-isolated nuclear relative to the oligo-dT selected poly(A)^+^ RNA pool could involve differences in RNA processing and turnover in the nuclear and cytoplasmic compartments. For example, differences in read coverage could reflect distinctions in the timing and fidelity of transcription, co-transcriptional processing, nuclear degradation, mRNA export to the cytoplasm, translation and cytoplasmic degradation. For example, the protein-coding transcripts enriched in the nucleus and depleted in the poly(A)^+^ pool could undergo destabilization following export to the cytoplasm, whereas those enriched in the poly(A)^+^ pool, which are presumed to be predominantly cytoplasmic, could be stabilized by active translation. Alternatively, these could be translationally inactive but stabilized in mRNA ribonucleoprotein complexes. Those transcripts with similar abundance in the two populations assayed, such as *BGAL2* and *WRKY68* (Fig. 5A, F), could be generally stable. The underlying reasons for the enrichment in nuclear pre-mRNA versus poly(A)^+^ transcript abundance could be explored by inhibiting transcription with the adenosine analog cordycepin or other compound and a subsequent time-course evaluation of the RNA in the two populations. Alternatively, mutants defective in post-transcriptional processes such as alternative splicing, cleavage / polyadenylation, mRNA export, nonsense-mediated decay, decapping-dependent decay or deadenylation-dependent decay could be combined with INTACT to explore specific post-transcriptional mechanisms.

## CONCLUSIONS

This study introduces the use of INTACT for the first time in a monocot, tests a new generation of NTF constructs, and evaluates nuclear transcriptome RNA-seq in a plant. The original version of INTACT developed for *Arabidopsis* and applied to tomato utilizes a protein domain of a RanGAP to place a biotinylated protein at the nuclear envelope by binding to an ONM protein (Deal and Henikoff, 2010; Ron et al., 2014). We directly transferred INTACT to rice using the rice RanGAP WPP and confirmed that it could be used for capture of nuclei. We also redesigned the INTACT construct to anchor the biotinylated protein to the ONM (Fig. 1). These constructs proved useful for isolation of nuclei from root and shoot tissue of seedlings grown in sterile culture (Fig. 2) and shoot tissue of greenhouse and field-grown plants. INTACT-obtained nuclei may have less contamination with ribosomes than nuclei isolated by conventional differential centrifugation (Fig. 3), which could prove beneficial for proteomic and other analyses. However, we cannot rule out the possibility that the reduced association of ribosomes with INTACT purified nuclei reflects methodological distinctions in capture of ONM associated rough endoplasmic reticulum.

Here we show that this method of nuclear isolation can be used to generate RNAseq libraries to monitor nuclear protein-encoding RNAs, following selective removal of rRNA. This subtraction of rRNA can be accomplished with commercial products, but also by the method demonstrated here that relies on hybridization of RNA to user-defined DNA oligos followed by a high-temperature double-stranded RNA digestion. We also determined the insertion site of the INTACT transgenes into the rice genome using an efficient method for T-DNA insertion site detection, establishing *OsNTF* genetic stocks for crossing to mutant genotypes.

The application of the INTACT method to root tips of rice enabled a comparison of the nuclear transcriptome to that routinely analyzed, steady-state poly(A)^+^ mRNA. This was accomplished by production of RNA-seq libraries with random hexamer priming of cDNA synthesis, the latter RNA captured by oligo-dT selection based on the attribute of a poly(A)^+^ tail. Deep sequencing of the libraries enabled a systematic comparison of the number, enrichment and RNA-seq read-distribution across individual transcripts. The results demonstrate that the two populations are not identical, as concluded from reassociation kinetic studies on tobacco nuclear and polysomal RNAs over 40 years ago (Goldberg et al., 1978). Of over 14,000 protein-coding gene transcripts detected in our study, >3000 were enriched significantly in the nuclear versus total poly(A)^+^ transcriptome, respectively. We did not monitor nuclear polyadenylated transcripts as their abundance is low and our goal was to assess the complete nuclear transcriptome. We hypothesize that nuclear-enriched mRNAs may be selectively nuclear-retained or unstable following their export to the cytoplasm, whereas poly(A)^+^- enriched mRNAs may be more rapidly synthesized, processed, and exported and maintained in the cytoplasm.

Access of nuclear contents enables monitoring of nuclear RNA, post-translational histone modifications associated with chromatin, DNA methylation and protein abundance, which can aid exploration of epigenetic and transcriptional regulatory mechanisms. INTACT can enable profiling all of these levels of regulation from the same preparation of nuclei, as shown for mouse neuronal cells (Mo et al., 2015). Access to nuclei can also improve chromatin immunopurification (Wang and Deal, 2015) and chromosome conformation studies (Rodriguez-Granados et al., 2016), enable RNA-protein interaction analyses (Foley et al., 2017), as well as refine proteomic analyses by allowing access to nuclei of discrete cell-or tissue-types (Amin et al., 2014). INTACT and other methods for isolation of nuclei vary in their advantages and disadvantages. INTACT has similar applications as FANS, used to assay nuclear poly(A)^+^ RNA, endoreduplication (Zhang et al., 2008), and regions of open chromatin by Assay for Transposase Accessible Chromatin-sequencing (ATAC-seq) (Lu et al., 2016; Bajic et al., 2017). INTACT constructs can be driven by near-constitutive, cell–type-and region-specific promoters, enabling assay of sub-population of nuclei of multicellular organs (Deal and Henikoff, 2010). Users can take advantage of the GFP signal from the NTF to define the expression patterns and confirm nuclear localization in cells of interest. As distinctions in nuclear and poly(A)^+^ mRNA populations (Fig. 5) could reflect distinctions in regulation of individual mRNAs, INTACT could be a good companion method for TRAP (Translating Ribosome Affinity Purification) that allows evaluation of RNAs engaged in translation (Zanetti et al., 2005; Mustroph et al., 2009). Such studies could advance understanding of the multiple levels of regulation that fine-tune the expression of individual genes. Moreover, use of the same promoters to drive INTACT and TRAP constructs might be used to target the same regions and cell types to obtain a comparative readout of nuclear pre-mRNA and translated mRNAs. The laser capture microdissection of tissue performed on rice (Jiao et al., 2009; Takehisa et al., 2012) and other species is not limited by genotype, whereas INTACT, fluorescent activated cell sorting and TRAP require transgenic plants (reviewed by (Bailey-Serres, 2013)). This disadvantage may be outweighed by the advantage of INTACT and TRAP over other methods of isolation of nuclei and transcripts from defined cells, as tissues may be cryopreserved prior to biochemical purifications. These technologies will likely complement single-cell transcriptomic and epigenomic analyses developed for animals but not yet reported in plants (Tanay and Regev, 2017). We propose that INTACT and TRAP are appropriate for evaluation of gene dynamics in individual cell types at multiple scales of regulation in response to environmental stimuli, which can help define genetic mechanisms critical to plant acclimation and adaptation to extremes in environment.

## MATERIALS AND METHODS

### Plant material, growth conditions and transformation

The rice genotype *O. sativa japonica* cv. Nipponbare was used for gene cloning and plant transformation. *Agrobacterium -* mediated transformation of embryogenic calli derived from rice mature seed embryos and plant regeneration were performed as described by Sallaud *et al*. After transfer to soil containing pots, plants were grown in a greenhouse (28°C day / 20°C night under natural light conditions) at the University of California, Riverside to obtain leaf tissue used for DNA analyses and the production of seeds. For isolation of nuclei and RNA, seeds from transgenic lines were dehulled and surface sterilized in 50% (v / v) bleach solution for 30 min and then rinsed with sterile distilled water. For nuclei isolation, seedlings were grown on plates (10 cm x 10 cm) containing Murashige and Skoog standard medium (MS) agar (1% w / v) and 1% w / v sucrose, during 7 days in a growth chamber (16 h day / 8 h night; at 28°C / 25°C day/night; 110 μEm^-2^s-^1^), in a greenhouse (28°C day / 20°C night under natural light conditions) or in the field during the autumn at the University of California, Riverside, Agricultural Experiment Station. The apical zone of the seminal and crown roots (1 cm root tips) or the shoot base was harvested directly into liquid nitrogen, pulverized and stored at-80° C until use.

### Nuclear tagging fusion (NTF) plasmids

A binary vector for INTACT in rice (*O. sativa* L.) was constructed as described in Ron *et al.*, 2014 using the Multisite Gateway vector pH7WG instead of pK7WG (https://gateway.psb.ugent.be/search). All primers used for construction are listed in Supplemental Table S3a. The NTF version 1 (*Os*NTF1) construct was made by replacing the *Arabidopsis thaliana* WPP domain for the one of rice (*OsWPP; LOC_Os05g46560.1*, AK242655) (Rose and Meier, 2001). This was comprised of a 345 nt region of the main open reading frame (ORF) (amino acids 1-115). These were introduced via partial digestion (*Stu*I / *Mfe*I) and ligation. The epitope-tagged *E. coli* biotin ligase (*mBirA* + *3xmyc*) was replaced by a synthesized (GenScript) codon use-optimized version (*Kpn*I / *Sna*BI), using the most frequently used codons in rice (http://www.kazusa.or.jp/codon/). For versions 2 and 3 (*Os*NTF2 and *Os*NTF3), the WPP domain was replaced with a C-terminal region of WIP to anchor the synthetic protein to the nuclear envelope. Enhanced GFP (eGFP) was amplified with primers to introduce *Stu*I / *Mfe*I sites at the 5’ and *Sna*BI / *Pac*I / *Avr*II at the 3’ end, respectively and incorporated to pTOPO-TA (Life technologies). The Biotin Ligase Receptor Peptide (BLRP) was amplified from the *Os*NTF1 binary vector to place *Stu*I / *Mfe*I sites at the 5’ and 3’ ends, respectively. This product was ligated upstream of eGFP. Inverted T-DNA *nos* and 35S CaMV terminators were amplified with *Sna*BI / *Pac*I sites and ligated downstream of eGFP. This cassette was digested and ligated to replace OsWPP–GFP-BLRP in the *Os*NTF1 binary vector using the sites *Sna*BI / *Stu*I. The correct orientation was checked by PCR. C-terminal domains from KASH protein coding regions *OsWIP2* (*LOC_Os09g30350.1*, AK241295.1) and *OsWIP3* (*LOC_Os08g38890.1*, AK071174) were amplified from root *Oryza sativa* L. cv. Nipponbare cDNA with primers to generate *Pac*I / *Avr*II sites (Xu et al., 2007). The PCR products were ligated to generate BLRPGFP-OsWIP2 (*Os*NTF2) and BLRP–GFP-OsWIP3 (*Os*NTF3) nuclear envelope-tagging proteins. For all binary constructs *Os*-codon BirA was made by coupling the promoter of *OsACT2* (*LOC_Os10g36650*) (from-2550 bp to the start codon, *Oryza sativa* L. cv. IR64) using a unique *Kpn*I site for cloning 3’ to the NTF and in the opposite orientation in the binary vector pH7WG backbone to generate the sequence verified constructs *p35S:OsNTF1 / OsWPP*, *p35S:OsNTF2/OsWIP2 and p35SOsNTF3 / OsWIP3*. Constructs were electroporated into *Agrobacterium tumefaciens* EHA105A prior to use in plant transformation.

### Generation of T-DNA enriched libraries for mapping of the insertion site

Young leaf tissue from transgenic lines was frozen in liquid nitrogen and stored at-80°C. For DNA extraction, 25 mg of sample was placed in a 2 mL micro-centrifuge tube along with three small stainless steel ball bearings. The samples were kept in liquid nitrogen before and after tissue disruption using a Tissue Lyser II (QIAGEN) for three 1 min cycles at 30 Hz, stopping every minute to chill the samples in liquid nitrogen. Tissue Lyser II blocks were pre-chilled in-20°C before use. Genomic (g) DNA was extracted as described in (Edwards et al., 1991), including two chloroform extractions. Extracted gDNA quality was assessed by electrophoresis in 1.2% (w / v) agarose gels and quantified using a Nanodrop ND-1000 UV-Vis spectrophotometer (Nanodrop Technology). When different samples contain regions near one T-DNA border that can be identifiable after sequencing, it is possible to pool DNA from different samples before library preparation. Pools of 1 μg gDNA were made and diluted to 100 μl with DNAse free water, then sheared by sonication (Bioruptor, Diagenode) in a 4° C water / ice bath at low intensity for 25 min with intervals set to 30 s on / 30 s off. The ice bath was replaced every 5 min. DNA fragments between 300 and 500 bp were selected using Agencourt AMPure XP beads according to the manufacturer’s instructions (Beckman Coulter). DNA binding to these beads depends upon the concentration of PEG-8000 in 1.25 M NaCl. Captured gDNAs were evaluated for the desired fragment size by electrophoresis in 2% (w / v) agarose gels.

Illumina-compatible libraries of 420-620 bp fragments were prepared by performing end repair, dA tailing, adapter ligation and amplification as described in Wang et al. (2011) with the following modifications. gDNA fragments were ligated with an universal adapter and amplified for 13 cycles of PCR using unique Tru-seq (Illumina) barcode adapters. To avoid the presence of adapter dimers, three rounds of AMPure XP bead purifications using 17.5% (w / v) PEG-8000 were performed, two before amplification and one after. Library quality was assessed using a Qubit^®^ fluorometer (Life Technologies), and the DNA 1000 Assay on an 2100 BioAnalyzer (Agilent Technologies). Library concentration ranged from 0.5-45 ng / μl and averaged 2.8 ng / μl, with and average size ∽500 bp. Based on concentrations obtained by DNA 1000 Assay, libraries were multiplexed to obtain a total of 500 ng and concentrated using a Speedvac concentrator (Savant).

Hybridization and sequence capture was achieved using a modification of the xGen^®^ Target Capture Protocol (IDT technologies). This method provides a means for enriching and sequencing specific regions of interest in a kit-less option to the procedures provided by Inagaki et al., 2015 and Jupe et al., 2014. Four 5’-biotinylated oligos targeting the right (RB) and left border (LB) sequences were used for the hybridization. Supplemental Table S3b lists primers used for target capture. Blocking oligos were designed to avoid cross hybridization of universal adapters. Salmon sperm DNA was substituted for C*ot*-human DNA. The hybridization mix was incubated at 62 °C for 48 h. DNA hybrids were captured for 30 min at room temperature using Streptavidin MyOne C1 Dynabeads (Life Technologies). Beads were washed for 5 min at 62 °C with 1X SSC (150 mM NaCl, 15 mM Na-Citrate pH 7.0), 0.1% (w / v) SDS and then with 0.1X SSC, 0.1% (w / v) SDS twice at 62 °C and once at room temperature. The final was was with wash with 0.2X SSC. After bead capture DNA was eluted by incubation for 10 min in 50 μl 125 nM NaOH, diluted with 50 μl Tris-HCl (pH 8.8) and purified using AMPure XP beads. T-DNA enriched DNA was and amplified by 14 cycles of PCR using primers P5 and P7. The quality of the final multiplexed libraries was assessed by use of the Qubit fluorometer and an Agilent Bioanalyzer DNA 1000 assay. The efficiency of hybridization and capture was quantified by quantitative real time PCR. Primers corresponding to the left and right T-DNA border sequences were used to quantify the abundance of insertion site fragments before and after hybridization and after final amplification. For a full bench protocol, see Supplementary Appendix I.

### Sequencing and data analysis for T-DNA insertion mapping

The Illumina-compatible library was sequenced on MiSeq (Illumina) using 500 cycles of MiSeq reagent kit v2 at the Institute for Integrative Genome Biology at the University of California, Riverside. The resulting 2 x 250 bp sequence data were analyzed using R statistical software. Paired-end reads containing overlapping sequences were merged to obtain longer reads using the FLASH (fast length adjustment of short reads; http://ccb.jhu.edu/software/FLASH/) module. The GREP (globally search a regular expression and print; http://www.gnu.org/software/grep/) module was used to filter out reads containing the left and right T-DNA border sequences, and the BOWTIE2 (http://bowtie-bio.sourceforge.net/bowtie2/index.shtml) module was used to map the border sequences against the rice genome (MSU Rice Genome Annotation Project Release 7). Regions of candidate insertions were identified using the chipseq module (http://www.bioconductor.org/packages/release/bioc/html/chipseq.html). See Supplementary Appendix I.

### Laser scanning confocal microscopy

Laser scanning confocal microscopy imaging was performed using a confocal Leica SP5. For images in Fig. 1 and S1, settings were 20X objective, 488 nm excitation laser at 50 % power, a 56.7 μm pinhole and 900 smart gain (SG), except in Fig. S1 for A1, G1: 800 SG, A2, G2: 650 SG. For images in Fig. 2, settings were 20X objective (zoom 10X), 405 nm excitation laser at 15% power, a 56.7 μm pinhole and 1050 SG.

### Nuclei purification by INTACT

Nuclei were purified from frozen and pulverized tissue as described previously for *Arabidopsis thaliana* (Wang and Deal, 2015) with only minor modifications including use of a 30 μm filter to exclude 30 to 70 μm cellular debris from the crude extract and extend centrifugation times. Tissue was resuspended in an ice-cold mortar containing 10 mL of freshly prepared nuclei purification buffer (NPB: 20 mM MOPS, 40 mM NaCl, 90 mM KCl, 2 mM EDTA, 0.5 mM EGTA, 0.5 mM spermidine, 0.2 mM spermine, pH, 7.0) containing 200 μL Protease Inhibitor Cocktail (0.4X, Sigma, P9599) per 50 mL of buffer. The homogenized extracts were filtered through a 30 μM nylon mesh (Celltrics) to remove cell debris and centrifuged at 1000 x g for 15 min at 4° C to pellet nuclei. Nuclei were resuspended in 1 mL of NPB (this suspension is considered input of the purification), 25 μL were kept for microscopic visualization and 200 μL for protein electrophoresis and western blot detection. Twenty-five microliters of M-280 streptavidin-coated Dynabeads (∽1.5 x 10^7^ beads; Life Technologies, catalog # 11205D) were added to the nuclei. This mixture was slowly rotated to allow capture of the nuclei in a cold room at 4° C for 30 min. The nuclei / beads suspension was diluted to 14 mL with NPB supplemented with 0.1% (v / v) Triton X-100 (NPB-T), in a 15 mL Falcon tube, mixed thoroughly and placed in a 15 mL magnet (adapted NEB 50 mL tube magnet) to capture bead-bound nuclei for 7 min at 4° C. The supernatant was carefully removed using a plastic Pasteur pipette, taking care to remove bubbles to avoid disturbing the beads. Beads were resuspended in 14 mL of cold NPB-T, placed on a rotating mixer for 30 sec, and then placed back in the 15 mL magnet to capture the beads-nuclei at 4° C for 7 min. This wash step was repeated and bead-bound nuclei were resuspended in 1 mL of NPB-T and transferred to a new tube. Estimation of nuclei yield and analysis of purity via protein detection and western blot was performed as described previously (Deal and Henikoff, 2011). For RNA extraction, the bead-bound nuclei mixture was placed on a 2 mL tube magnet for 4 min at 4° C to capture the nuclei, which were resuspended in 20 μl NPB before proceeding to the extraction. For comparison, nuclei were isolated without the aid of INTACT according to the protocol described by (Choudhary et al., 2009).

### Protein detection by western blotting

Crude cellular protein, input of INTACT and purified nuclei were separated by 12% (w / v) SDS-PAGE and silver-stained or used in immunoblot analyses. Proteins loaded in Figure 3 panels B, C and D were quantified using Qubit Protein Assay Kit. This dye-based method overestimated the concentration of nuclear samples because of their higher concentration of proteins with elevated pI such as histones. Biotinylated proteins were detected using horseradish peroxidase (HRP)-conjugated streptavidin (1:2000; ThermoFisher S911). Histone H3 was immunodetected using rabbit polyclonal antiserum (1:1,000; Abcam 1791). The small ribosomal subunit protein RPS6 was immunodetected using a rabbit polyclonal antiserum against *Zea mays* RPS6 (1:5,000; Williams et al., 2003) as primary antibody and an HRP-conjugated goat anti-rabbit IgG as secondary antibody (1:10,000; Bio-Rad). Visualization was by use of a chemiluminescence reagent according to the manufacturer’s instructions (Luminata Crescendo Western HRP Substrate, Millipore, WBLUR0500) followed by exposure to x-ray film (Bioland, A03-02). Molecular mass markers (Pageruler 10-180 kDa, Thermo-Scientific) were used for each run.

### Total and nuclear RNA extraction, rRNA degradation and short-read library preparation

Total poly(A)^+^ RNA was extracted from frozen tissue using polysome extraction buffer (Mustroph et al., 2009), followed by direct capture of poly(A)^+^ RNA using a biotinylated oligo dT according to Townsley et al., 2014. Nuclear RNA was extracted from 66,000-117,000 (Supplemental Table S2a) INTACT-purified nuclei using the RNeasy Micro kit (Qiagen), DNase-treated with Turbo DNase I (ThermoFisher Scientific) and processed with Agencourt RNAClean XP beads (Beckman Coulter), per the manufacturer’s’ instructions. Ribosomal RNA (rRNA) was removed from the RNA using degradation method described by (Morlan et al., 2012) which employs an RNase that specifically degrades RNA species in RNA:DNA duplexes, targeted by use of DNA oligomers that correspond to precursor and mature rRNA encoded in the nucleus.

To accomplish removal of specific abundant rRNA, 60 bp DNA probes were designed and synthesized (Integrated DNA Technologies) to complement the rice (*O. sativa)* nuclear rRNA sequences and their transcribed spacers. A maximum of six mismatches per probe was allowed for targeting *O. sativa, Medicago truncatula* and *S. lycopersicum*, using additional species-specific probes when common probes were not possible due to sequence variation. Rice probes are listed in (Supplemental Table S3c) and were based on GenBank accessions KM036282 (18S rRNA-5.8S rRNA-26S rRNA including the complete internal transcribed spacer (ITS) and external transcribed spacer (ETS) regions) and DQ152232.1 (5S rRNA).

For degradation, probes were hybridized with up to 1 μg RNA in a 6 μl reaction containing 0.1 μM of each probe and 1 μl of 5x hybridization buffer (0.5 M Tris-HCl [pH 7.0], 1 M NaCl). The probe concentration was scaled down proportionally for lower amounts of RNA. For denaturation and hybridization, the sample was heated to 95 °C for 2 min, cooled to 45 °C at 0.1 °C / s and incubated at 45 °C for 5 min. For RNase treatment, the hybridization reaction at 45 °C was increased to 10 μl by adding 5 U of thermostable RNase H Hybridase (Epicentre) and 1 μl of Hybridase buffer (500 mM Tris-HCl [pH 7.4], 1 M NaCl, 200 mM MgCl_2_), and incubated at 45 °C for 30 min. To remove the DNA probes and degraded RNA from the samples, the sample was re-treated with Turbo DNase I (ThermoFisher Scientific) and cleaned up with Agencourt RNAClean XP beads (Beckman Coulter). For a full bench protocol, see Supplementary Appendix II. RNA-seq libraries of pre / rRNA subtracted nuclear and poly(A)^+^ selected total mRNA were prepared as described by Townsley et al. 2014.

### RNA-seq and data analysis

Total poly(A)^+^ and nuclear RNA libraries were sequenced on HiSeq 3000 to obtain 50 nt single-end reads. All data analysis steps were performed on the University of California Riverside Institute for Integrative Genome Biology high performance bioinformatics cluster (http://www.bioinformatics.ucr.edu/), supported by NSF MRI DBI 1429826 and NIH S10-OD016290. R packages from Bioconductor including systemPipeR (Girke, 2014) were used. Quality reports of raw reads were generated with the FastQC package (http://www.bioinformatics.babraham.ac.uk/projects/fastqc/). Raw reads were mapped with the splice junction aware short read alignment suite Bowtie2 / Tophat2 to the mitochondrial and chloroplast genome as a first filter before mapping to IGRSP1.0-30 rice sequence (http://plants.ensembl.org/Oryza_sativa/Info/Index), allowing unique alignments with ≤2 nt mismatches. Expression analyses were performed by generating read count data for features of exons–by-genes using the summarizeOverlaps function from the GenomicRanges package (Juntawong et al., 2014). Statistical analysis of differentially expressed genes and other feature types was performed with the Generalized Linear Model approach from the EdgeR package (http://bioconductor.org/packages/release/bioc/html/edgeR.html) to obtain fold change (fold difference) and false discovery rates (FDRs) confidence filters, applied as indicated. To increase robustness only genes identified as differentially expressed with both DEseq2 and EdgeR were used for downstream analysis. Enrichment analysis of Gene Ontology (GO) terms was performed with systemPipeR using the BioMart database. Gene feature read counts were determined over ranges produced with the function genFeatures on systemPipeR. Coverage plot over transcripts were calculated and produced with the functions computeMatrix and plotHeatmap from deepTools (http://deeptools.readthedocs.io/en/latest/index.html). Genes containing a high intron / exon coverage ratio were visually inspected using IGV and filtered from the analysis of feature counting and coverage plots, as these were high abundance short sequences that spuriously mapped to introns (Supplemental Table S4). Short read sequence data were deposited in the Gene Expression Omnibus database (accession no. GSE99122).

### Data visualisation and clustering

BAM files were generated for each sample. These were processed into bedgraphs using to bedtools genomecov command (Quinlan and Hall, 2010) and scaled by number of aligned reads to exonic regions of genes in each bam using the-*scale* argument. Bedgraph files were compressed to bigwigs using *bedGraphToBigWig* (http://genome.cse.ucsc.edu/index.html) and visualized using the Integrative Genome Browser (http://software.broadinstitute.org/software/igv/). Scatter plots and density plots were made using the *ggplot2* (Wickham) library for R.

## ACKNOWLEDGEMENTS

We thank Maureen Hummel, Michael Covington, Marko Bajic, Donnelly West and Kristina Zumstein for active discussions throughout this project.

## Supplemental Data

Supplemental Figure S1. NTFs subcellular localization for a second independent transformation event.

Supplemental Figure S2. Detection of biotinylated *Os*NTF proteins in rice.

Supplemental Figure S3. Nuclei isolation from roots using INTACT from a second independent event.

Supplemental Figure S4. Coverage of gene features for nuclear and poly(A)^+^ RNA libraries.

Supplemental Figure S5. Complexity evaluation of nuclear and poly(A)^+^ RNA libraries.

Supplemental Table S1. Transgenic lines insertion sites.

Supplemental Table S2. Nuclear RNA and total poly(A)^+^ transcriptome dataset.

Supplemental Table S3. Primer and probe lists.

Supplemental Table S4. List of transcripts with high intron/exon coverage ratio

## Supplemental Methods

Appendix I. Mapping of T-DNA insertion sites using sequence capture and IlluminaMiseq.

Appendix II. Method for rRNA degradation.

**Supplemental Figure S1: NTFs subcellular localization for a second independent transformation event.**

Representative confocal images of GFP fluorescence in organs from 7–d-old plants grown in a growth chamber expressing a *p35S:OsNTFs*. **A1** to **F**: root MZ; **G1** to **I**: root DZ, 1 cm from the root tip, epidermal cells; **J** to **L**: blade region between the midvein and outer margin of a newly expanding leaf blade, guard cells and epidermal cells are visible. Confocal microscope settings are same as Fig. 1 (see Methods) for *p35S:OsNTF2* and *p35S:OsNTF3*. For *p35S:OsNTF1*, the second independent line (**A1** and **G1**) selected for this study showed a strong expression of the NTF. Leaves are maximum intensity Z-projections of a confocal Z series. Scale bars are: **A1-C**, **G1-L**: 25 μM, **D-F**: 5 μM.

**Supplemental Figure S2: Detection of biotinylated OsNTF proteins in rice.**

Total proteins were extracted from 7–d-old roots of seedlings in 2X SDS buffer and fractionated by electrophoresis in an 12% (w / v) polyacrylamide-SDS gel, transferred to a membrane and processed with HRP-conjugated streptavidin and commercial monoclonal antibody against Histone 3 (H3) (α-Histone H3) as a loading control. Non-transgenic control (Nb) shows the presence of an endogenous biotinylated product (*), a red arrow indicates proteins corresponding to *Os*NTF2 and 3, whereas a blue arrow indicates *Os*NTF1. Molecular weight markers are shown on the left. The expected molecular weight of Histone H3, *Os*NTF1, 2 and 3 are 14 kDa, 43.09 kDa, 52.87 kDa and 52.91 kDa, respectively.

**Supplemental Figure S3: Nuclei isolation from roots using INTACT from a second independent event**

**A,** Transgenic lines carrying *35Spro-OsNTF1*, *2* or *3* and *ACT2pro-BirA* shown in Fig. S1 and a non-transgenic control (Nb) were used as starting material. Each line is distinct from that in Fig. 2. Nuclei were isolated from roots of 7–d-old seedlings by use of streptavidin-coated magnetic beads, stained with propidium iodide for visualization and counted by use of an hemocytometer. **B,** Detection of Histone 3 (H3) protein in nuclei before (Input) or after purification by INTACT (INTACT nuclei). For each sample, an equal amount of starting material (0.5 g) was used to obtain nuclei and a proportional amount of each yield was solubilized in an SDS-containing buffer and fractionated by electrophoresis in an 12% (w / v) polyacrylamide-SDS gel, transferred to a membrane and processed with commercial monoclonal antibody against Histone 3 (H3) (α-Histone H3). Molecular weight marker is shown on the left. The expected size of Histone H3 is 15.27 kDa.

**Supplemental Figure S4. Complexity evaluation of nuclear and poly(A)^+^ RNA libraries.**

Each library was randomly subsampled for a specific quantity of reads to evaluate the number of individual genes detected with a specific number of reads counts. **A,** Genes detected with using a minimum threshold of > 5 rpkM in four nuclear and poly(A)^+^ bioreplicates with increasing coverage. **B,** Genes detected above a high minimum threshold (> 10 rpkM).

**Supplemental Figure S5: Coverage of reads on gene features for nuclear and poly(A)^+^ libraries.**

Aligned reads were binned into groups of features corresponding to 5’ and 3’ untranslated regions (5’ UTR, 3’UTR), introns, and coding sequences (CDS). An average of 29.46 million poly(A)^+^ and 1.74 million nuclear RNA reads, respectively, were mapped per replicate among these features. For nuclear RNA, highly abundant reads on introns with no correlation to exon reads for the corresponding transcript were excluded from the analysis. The *x*-axis indicates the bioreplicate number for each sample and the *y -* axis indicate the proportion of the total reads mapped to a specific feature.

### SUPPLEMENTAL RESULTS

#### Development of an Inexpensive Method for Mapping T-DNA Insertion Sites

To determine the number and the site of transgene insertions we sequenced T-DNA enriched libraries, obtained by hybridizing multiplexed and barcoded libraries of fragmented genomic DNA to biotinylated probes corresponding to the T-DNA right (RB) and left (LB) sequences and subsequent magnetic capture. High-throughput sequencing of 250 nt from both ends enabled identification of the genomic region flanking individual insertions. The capture efficiency prior to amplification before sequencing was 19.8% for the LB and 13.8%, for RB regions, as compared to 0.003% for a non-targeted region. After PCR amplification the LB and RB sequences were enriched on average ∽ 180-and ∽ 186-fold relative to the input. By contrast, there was no *UBCII* enrichment (<0.003-fold). The final multiplexed, T-DNA border-enriched and amplified library yielded 154,342 reads with RB or LB sequences flanking rice genomic sequences. Of the 12 transgenics evaluated, 17 T-DNA insertion locations were mapped. Transgene insertion was independently confirmed for 6 lines by design of primers that enabled amplification from the T-DNA to the genomic bordering sequences (Supplemental Table S1).

